# The enlightened entomologist: fast, non-destructive whole-arthropod clearing for three-dimensional imaging

**DOI:** 10.64898/2026.09.02.748800

**Authors:** Marwa Moulzir, Hugo Touja, Martin Oheim, Brigitte Delhomme

## Abstract

Arthropods are a strikingly diverse phylum of invertebrates with segmented bodies, chitinous exoskeletons, and jointed limbs, many of which are colloquially referred to as insects. Their pigmented and optically dense bodies pose a considerable challenge for microscopy. While early naturalists focused on external features, modern approaches combining tissue clearing and fluorescence microscopy seek to explore internal anatomy in three dimensions. However, both chemical clearing and three-dimensional (3-D) microscopy are specialized techniques and often require considerable adaptation and optimization across species. Here, we introduce a versatile, fast, effective and non-toxic clearing method that renders diverse arthropods transparent within hours to days. Existing upright microscopes or macroscopes can be upgraded with a modular and affordable light-sheet microscope, allowing rapid volumetric imaging of arthropods. Our pipeline is fully compatible with dye staining and immunofluorescence labeling, while endogenous autofluorescence provides valuable anatomical context and facilitates 3-D reconstruction.

**Disclaimer:** The authors of this article are not specialists in entomology. Accordingly, this work is not intended to provide a taxonomic or entomological reference description, but rather to offer specialists of the domain a new, rapid workflow that allows examining the detailed 3-D anatomy and thus facilitating their studies.

## Introduction

More than two-thirds, and possibly up to 90%, of all known animal species are arthropods [1], many of which are commonly referred to as “insects” by non-entomologists. With their characteristic segmented body structure, arthropods have long captivated humans through their remarkable diversity and beauty. Their metamorphosis during development has triggered fascination, embedding entomology, the scientific study of arthropods, deeply within human history. Easy to collect and gratifying to observe, arthropods, with their bewildering range of sizes, shapes, colors and textures, became a favored subject of early microscopists and taxonomists. They fill the collections and displays of natural history museums and they continue to fuel the enthusiasm of amateur collectors and naturalists [2]. This enduring fascination traces a line from pre-enlightenment cabinets of curiosities and ornate specimens pinned beneath glass to contemporary tourist attractions like the Dubai Butterfly Garden or its more modest French counterpart, L’île aux Papillons.

Humans have long relied on arthropods for pollination and ecosystem services, but they can also cause harm, such as locust swarms that damage crops in North Africa, an issue intensified by climate change. Meanwhile, saproxylic arthropods are essential for decomposing deadwood, especially in warm tropical and subtropical regions [3]. More surprisingly, certain arthropods and their larvae also possess the ability to degrade plastics through specialized gut enzymes and symbiotic microbiota [4].

From a nutritional standpoint, arthropods are a sustainable source of protein, essential amino acids and micronutrients. Insect powders are being used in animal food and edible arthropods have long been part of human diets [5,6]. Honey is a natural sweetener and carmine dye, from cochineal (*Dactylopius coccus*), is used in food, pharmaceuticals and cosmetics. Lac and shellac, resinous secretions from arthropods like the lac bug (*Kerria lacca*), are key components in polishes and cosmetics like hairspray, eyeliner and mascara. Sericin, a protein secreted by silkworm larvae (*Bombyx mori*), is incorporated into cosmetic formulations [7]. On the other hand, arthropods also transmit major diseases: mosquitoes spread malaria, dengue fever and Zika; ticks carry Lyme disease bacteria; and scorpions, bees, spiders and wasps can cause medically significant envenomation [8,9]. Even after death, arthropods remain linked to humans, as insects colonize corpses in predictable waves used in forensic entomology to estimate post-mortem interval [10].

Thus, while arthropods play critical roles in ecosystems, human nutrition, industry and medicine, they can also cause major agricultural damage and transmit serious diseases. Studying insects remains important as it allows the detailed observation of their anatomy and biology which helps scientists better understand their ecological functions, health impacts and potential applications.

Owing to these diverse roles and complex morphologies, there has long been a quest for an accurate volume representation of arthropods to better study and preserve their structure. Early scientific illustrators struggled to convey the three-dimensionality of arthropods in flat drawings, an issue that has been addressed with modern volume-imaging techniques including brightfield [11] confocal [12], two-photon microscopy [13] or electron microscopy [14]. More recently, light-sheet microscopy has shown promise for studying arthropods, such as insect embryos [15,16] and optically cleared adult Drosophila specimens [17,18], but applications have remained largely restricted to a limited number of species and protocols.

Full 3-D imaging of the arthropod body has remained challenging because its rigid, pigmented cuticle (rich in chitin and melanin) is opaque and often reflective, creating iridescence through structural color [19]. Refractive index (RI) differences among internal tissues additionally scatter light. A common strategy for facilitating volume imaging is chemical tissue clearing [20,21] which makes tissues transparent by reducing light scattering through lipid removal and RI matching. In arthropods, it allows the visualization of internal structures (e.g., nervous system, gut, muscles) and is also valuable for small insects like ants that are challenging to dissect. However, despite progress, tissue clearing in arthropods suffers from many limitations: species and tissues respond unevenly, strong pigmentation can still block light, and generally protocols are time-consuming and labor-intensive. Additionally, some methods require toxic chemicals and specialized lab equipment, while clearing agents may also alter fluorescence and render tissue brittle, which adds another hurdle to imaging [22].

In the current work, we introduce a simple, fast and non-toxic clearing workflow adapted to both confocal and light-sheet microscopy for 3-D arthropod imaging. This allows label-free 3-D whole-arthropod visualization in great detail, alone or paired with histological stains and immunofluorescence.

## Results

### Hydrogen peroxide-based solution bleaches most arthropods

The depigmentation solution used effectively bleached a wide range of arthropods with diverse pigment compositions: melanin in spiders, xanthopterin in wasps and hornets [23], erythropterin in firebugs and harlequin bugs. Incubation times varied greatly according to the species, ranging from 4 h for small specimens like woodlice to 17 days for larger ones (Table 1) (Fig. 2C). Of note, the duration of this step can be significantly reduced when incubation is performed at 37 °C and under moderate agitation. For example, depigmentation of a hornet required approximately 55 days at room temperature without agitation, whereas the addition of heat and agitation reduced the incubation time to about 8 days. Regular replacement of the depigmentation medium also substantially accelerates the process, as freshly prepared solutions maintain higher efficiency.

**Table 1:**
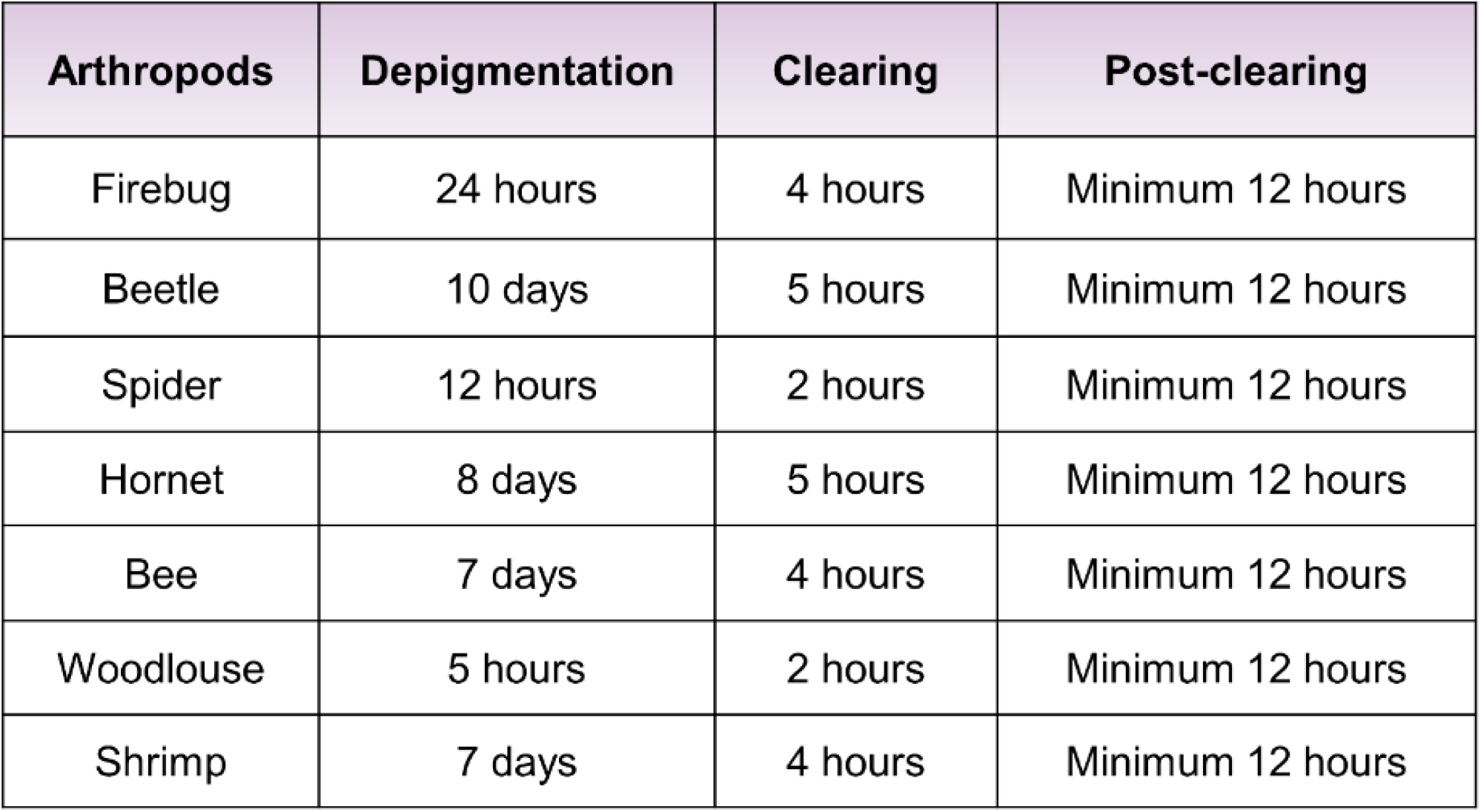
Typical time scales for arthropod depigmentation and clearing. Summary of the time required for each stage of the clearing protocol for different arthropods (firebug, beetle, spider, hornet, bee, woodlouse and marine shrimp).

| Arthropods | Depigmentation | Clearing | Post-clearing |
| --- | --- | --- | --- |
| Firebug | 24 hours | 4 hours | Minimum 12 hours |
| Beetle | 10 days | 5 hours | Minimum 12 hours |
| Spider | 12 hours | 2 hours | Minimum 12 hours |
| Hornet | 8 days | 5 hours | Minimum 12 hours |
| Bee | 7 days | 4 hours | Minimum 12 hours |
| Woodlouse | 5 hours | 2 hours | Minimum 12 hours |
| Shrimp | 7 days | 4 hours | Minimum 12 hours |

While most of the coloration was lost (Fig. 1, middle column), some species showed a residual brown tint on the cuticle. The remaining hue does not seem to adversely affect image acquisition or the resolution. One explanation might be that H_2_O_2_ effectively bleaches melanin, but not UV-protective tyrosine derivatives present in some arthropods such as European rhinoceros beetle (*Oryctes nasicornis*), a large black coleopteran measuring approximately 3 cm × 2 cm × 2.5 cm, even after 17 days of depigmentation (Fig. 2C).

**Figure 1:**
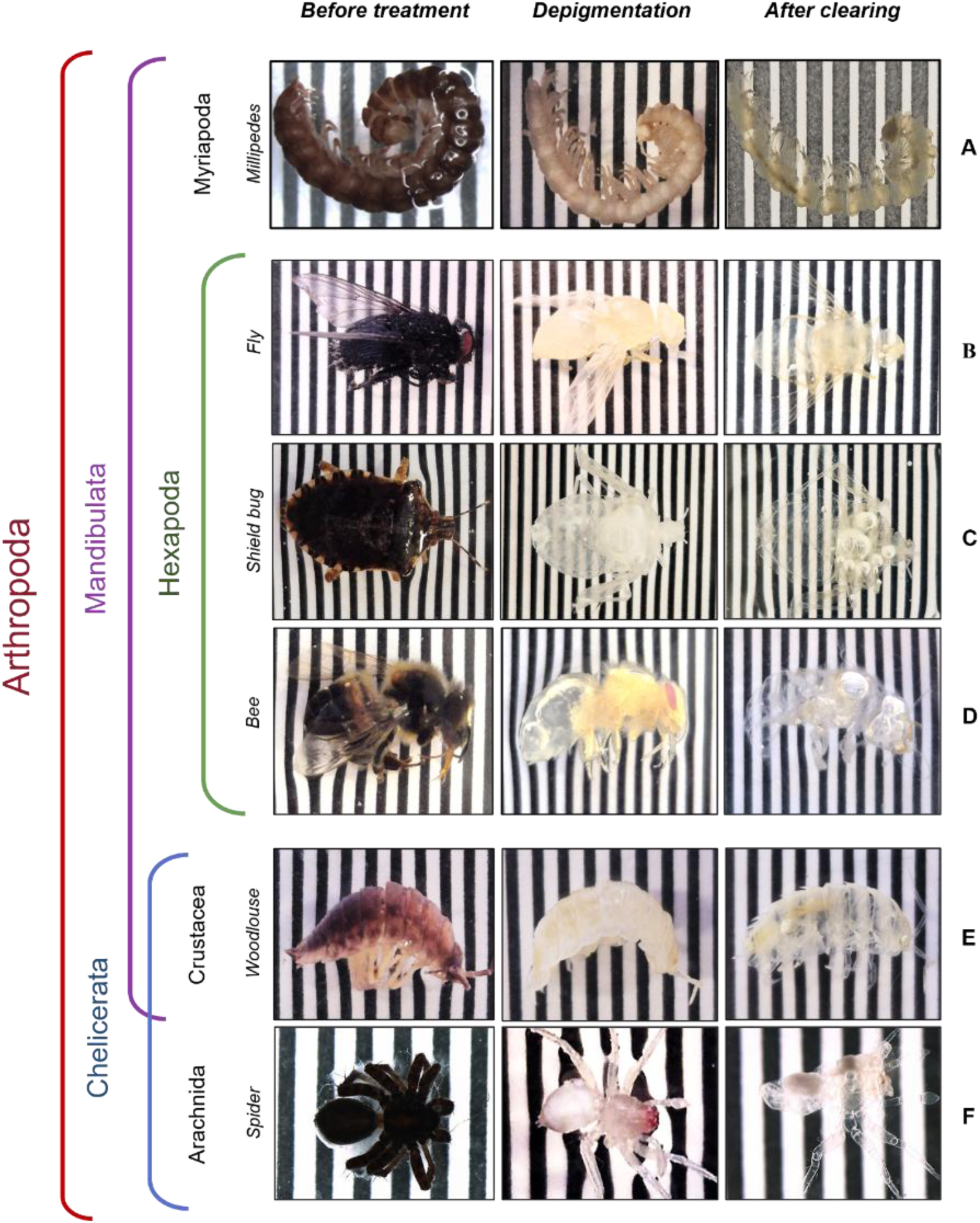
Clearing of various arthropod specimens. Representative specimens of different arthropod classes and subphyla shown at three stages of the protocol: fixed (*far left column*), depigmented (*middle column*) and cleared (*far right column*). From top to bottom: Diplopoda *(**A**, millipede)*, Hexapoda *(**B–D**, insects)*, Crustacea *(**E**, woodlouse)* and Arachnida *(**F**, spider)* (Pycnogonida and Merostomata not included). Scale bar: black/white stripe width: 1 mm.

**Figure 2:**
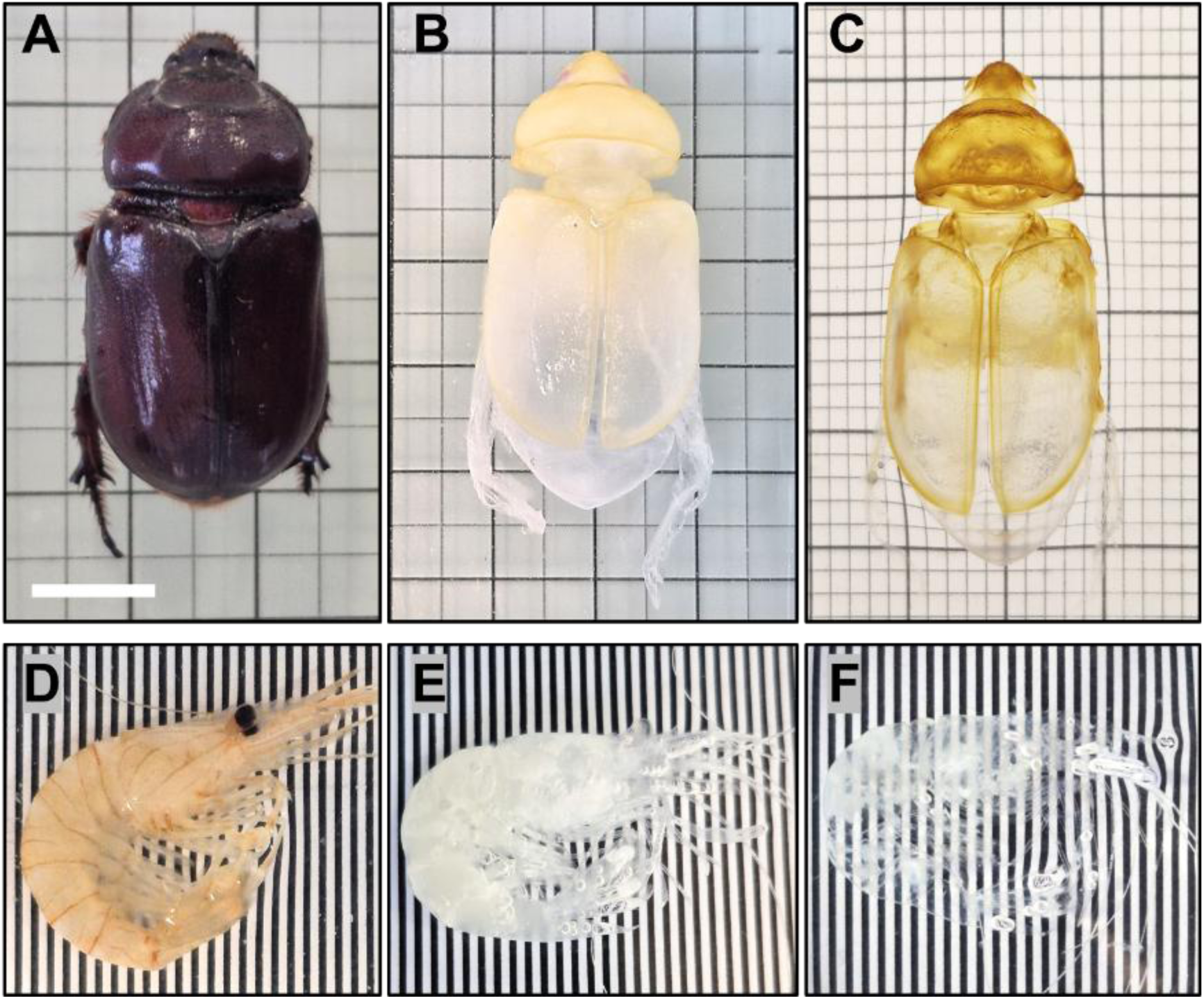
Clearing of large arthropods. Female European rhinoceros beetle *(Oryctes nasicornis)* and marine shrimp at different clearing stages. (***A****, **D***) before treatment, (***B****, **E***) after depigmentation and (***C****, **F***) after clearing, respectively. The level of transparency achieved makes the background grid clearly visible through the specimens. Scale bars: (***A***) 1 cm, (***D-F***) black/white stripe width 1 mm.

Also, some arthropod’s eyes, like the spider’s, were particularly resistant to depigmentation and retained a red coloration; however, it disappeared by the end of the clearing process which resulted in highly transparent specimens (Fig. 1F). Finally, we noted in some species bubble formation, a process particularly pronounced in marine shrimp (Fig. 2F).

### Detailed whole-arthropod structural imaging via clearing

Fig. 1 illustrates the successful clearing across multiple arthropod classes, including *Myriapoda* (millipedes), *Arachnida* (spiders), *Insecta*, and *Crustacea*, the latter represented by a terrestrial woodlouse (Fig. 1E) and a marine shrimp (Fig. 2F). Clearing was achieved using the same cocktails regardless of arthropod size.

Maximum transparency was typically achieved within 2-48 h for all species and reached, for example,70% transparency for the harlequin bug (Fig. 6B), 80% for the wasp (Fig. 4A, bottom), 70% for the spider, and 80% for the shield bug (Fig. 1C). However, quantifying the transparency of 3-D samples is inherently challenging due to the presence of multiple overlapping tissue layers. Consequently, 2-D images often fail to accurately reflect the true degree of transparency observed visually. The European rhinoceros beetle (*Oryctes nasicornis)* required 5 days for delipidation but achieved 60% transparency within 48 h after incubation in the post-clearing solution (Fig. 2A–C).

**Figure 3:**
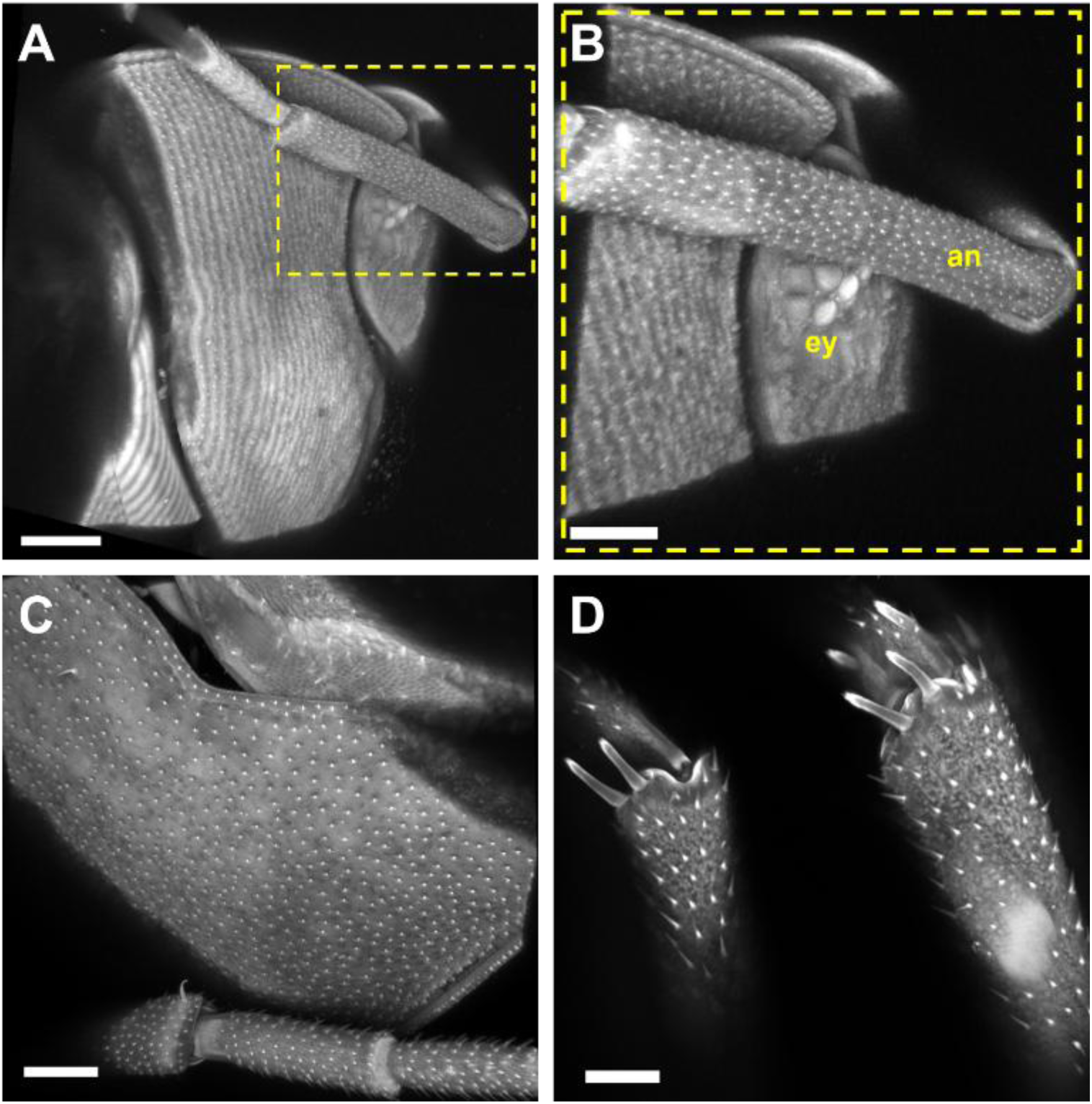
Confocal autofluorescence imaging. Label-free confocal imaging of woodlouse *(Porcellio scaber)* with a 10×/NA 0.45 objective excited upon 405 nm. *(**A**)* AF z-projection of the head region, scale bar: 400µm. *(**B**)*. Insets: crop view of compound eye and antenna, respectively, scale bar: 200µm. *(**C**)*. AF z-projection of exoskeleton, scale bar: 200µm. *(**D**)*. AF image of the legs’ articulations, scale bar: 100µm. ***Abbreviations:*** ey-eye; an-antenna.

**Figure 4:**
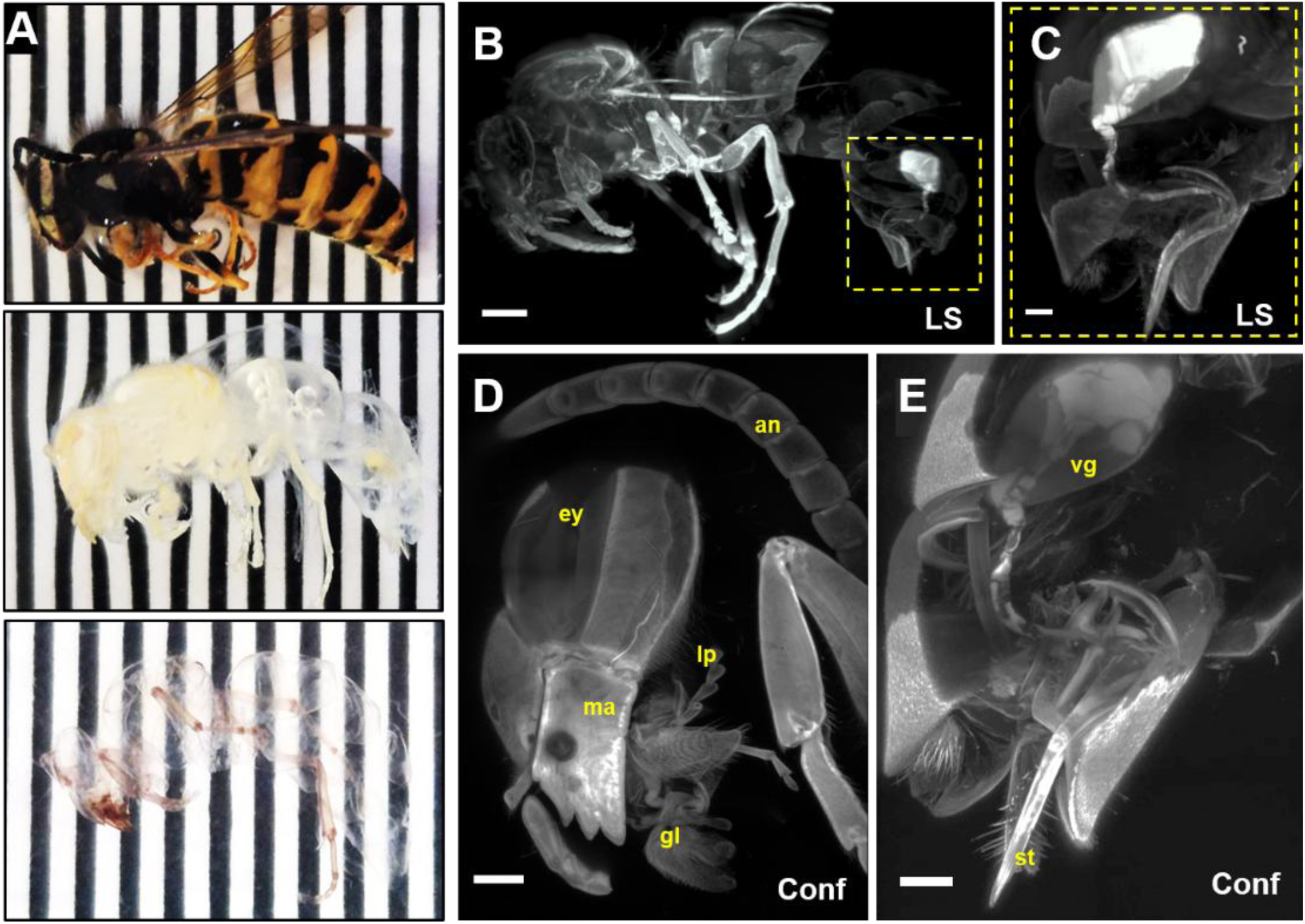
Confocal and light-sheet autofluorescence imaging of a cleared wasp. *(**A**)* Wasp before treatment *(top),* after depigmentation *(middle)* and after clearing *(bottom)*, scale bar: black/white stripe width 1 mm. *(**B**)* Light-sheet AF z-projection of the whole wasp body *(ex 488 nm)*, scale bar: 2mm. *(**C**)* Inset: magnified view of the venom gland and stinger *(ex 488 nm, acquisition time 600ms per image frame),* scale bar: 300 µm. *(**D**)* Confocal AF z-projection of the wasp head with a 10×/NA 0.45 objective *(ex 405 nm)*, scale bar: 500 µm. *(**E**)* Confocal AF z-projection of the venom gland and stinger with a 10×/NA 0.45 objective *(ex 405 nm, acquisition time of 31s per image frame),* scale bar: 300 µm. ***Abbreviations:*** LS-light-sheet microscope; Conf-confocal microscope; vg-venom gland; st-stinger; ma-mandible; gl-glossa; lp-labial palpus; ey-eye; an-antenna.

**Figure 5:**
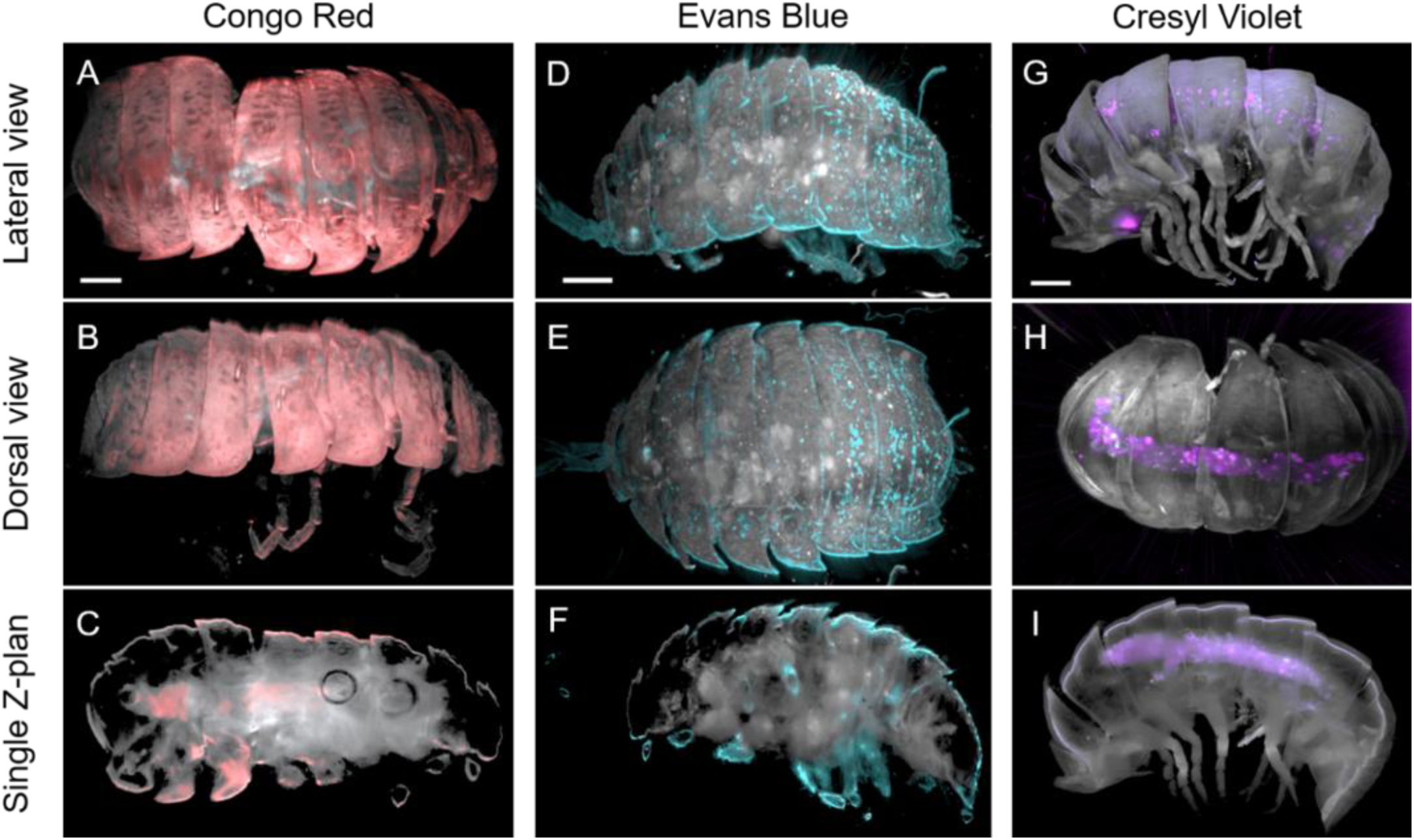
Whole-arthropod staining and 3-D imaging of cleared woodlouse. Cleared woodlice stained with Congo red (*left column*), Evans blue (*middle*) and Cresyl violet (*right*). Images are 3-D renderings of light-sheet microscopy z-stacks acquired using 488-nm excitation for AF *(grey)* combined with dye-specific excitation/emission settings *(Congo red, 561nm, red), (Evans blue, 638nm, blue), (Cresyl violet, 638nm, purple)*. 3-D reconstructions were generated using Imaris Viewer. The upper panels *(**A, D, G**)* show lateral views of the reconstructed specimens, the middle panels *(**B, E, H**)* dorsal views and the lower panels *(**C, F, I**)* show single optical z-planes extracted from the z stacks. Congo red and Evans blue mainly label the exoskeleton, while Cresyl violet labels internal tissues. Scale bars: Congo red and Evans blue 1 mm, Cresyl violet 2 mm.

**Figure 6:**
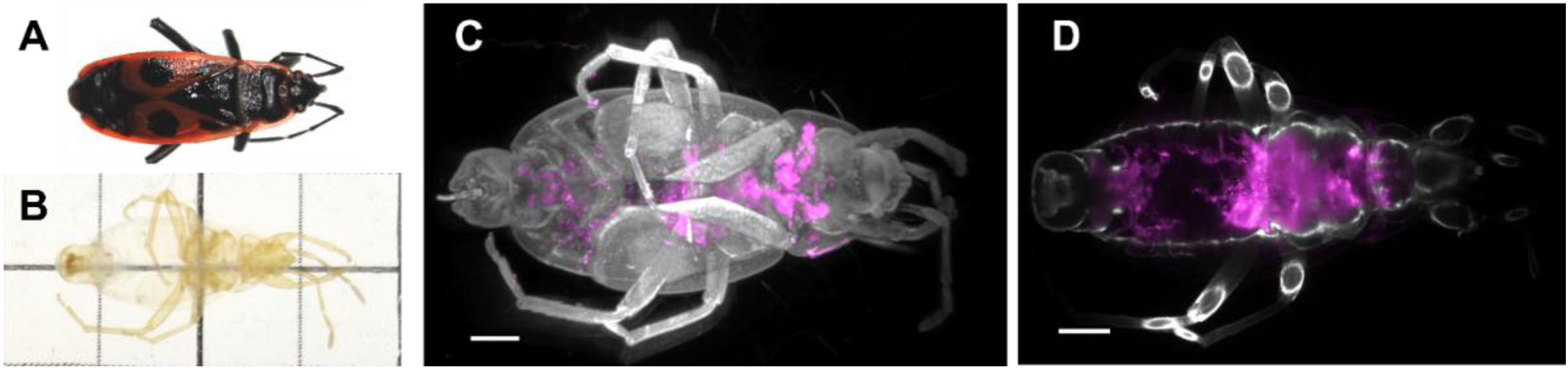
Immunolabeling of a cleared firebug. *(**A**)* Specimen before clearing. *(**B**)* Same specimen after clearing. *(**C**)* Light-sheet 3-D reconstruction showing tissue autofluorescence *(488nm, grey)* and Tuj1 immunolabeling *(561nm, magenta) (**D**)* Single optical z-plane showing autofluorescence *(grey)* and Tuj1 labeling *(magenta)*. Scale bar: 200 µm.

### Comprehensive 3-D visualization of internal anatomy using label-free imaging

Insect autofluorescence (AF) mainly originates from molecules associated with the cuticle and from naturally fluorescent cellular cofactors [24]. The strongest signal typically arises from cuticular sclerotization products, generated during exoskeleton hardening when catecholamines such as N-acetyldopamine (NADA) and N-β-alanyldopamine (NBAD), cross-link with cuticular proteins and chitin. These reactions produce conjugated structures that fluoresce strongly under UV–blue excitation. Resilin, an elastic protein present in joints and wing hinges, also contributes intense fluorescence due to cross-links formed by di- and tri-tyrosine residues [25]. Additional AF signals can originate from eye pigments, particularly from fluorescent pteridines, and from common intracellular cofactors such as NADH/NADPH and flavin cofactors (FAD, FMN) found in metabolically active tissues. Together, these compounds generate a spectrally broad AF that can be exploited for label-free imaging of arthropod anatomy.

We therefore investigated whether label-free 3-D imaging alone could provide sufficient contrast following tissue clearing. To better characterize arthropod AF, we imaged a harlequin bug (*Graphosoma lineatum*) before and after clearing at excitation wavelengths of 488, 520, 658 and 784 nm (Supplementary Fig.1). The highest signal and greatest detail were observed upon 488-nm excitation. In contrast, only minimal AF was detected in the near-infrared (NIR) range, providing a spectral window to reduce background signal and enhance contrast during targeted immunostaining with NIR-emitting secondary antibodies.

We next imaged a variety of arthropods, either using a confocal laser-scanning microscope or a custom-built light-sheet microscope. The overall picture emerging from our multi-wavelength excitation/emission imaging is that the emission filter has limited impact provided that it collects broad AF emission, and that most of the fluorescence is excited at 405 nm or 488 nm, with minor contributions from AF excited at 561 and 633 nm.

Confocal microscopy (405-nm excitation) enabled high-resolution 3-D AF imaging of a woodlouse’s anatomical features such as legs, exoskeleton, eyes and antennae (Fig.3). It also allowed us a detailed visualization of a wasp head [26] (Fig. 4). Acquisition of a z-stack allowed imaging to depths of up to 1300 µm before significant signal attenuation. Due to the sequential point scanning, confocal microscopy scans the sample point by point with a focused laser which produces high spatial resolution optical sections but is limited by long acquisition times for large-volume datasets. Light-sheet microscopy illuminates the sample with a thin plane of light which enables rapid imaging of large volumes and reduces phototoxicity and photobleaching compared to confocal microscopy. Therefore, it offers a quicker alternative, as illustrated by comparative imaging of the wasp abdomen, where structures such as the venom gland and stinger are clearly resolved (Fig. 4C–E).

Light-sheet AF imaging (488 nm excitation) showed strong fluorescence in the venom glands and weaker signal in the stinger. Conversely, confocal imaging at 405 nm reduced gland fluorescence while enhancing the stinger signal, highlighting the importance of optimizing the excitation wavelength for structure-specific visualization. In all examined arthropods, the articulations consistently exhibited strong AF signals. Also, the wasp’s abdomen with its black and yellow stripes, which came optically transparent after clearing, showed a distinct pattern: the melanin-rich regions (originally black) emitted very weak AF upon 488-nm excitation, whereas the initially yellow pigmented regions consistently emitted a more intense AF signal. This difference can serve as a reference point for 3-D navigation within the sample (Fig. 4B).

### Combined dye and label-free imaging improves structural context

As previously observed in mammalian tissues (Delhomme et al., unpublished), the nuclear dye DAPI, as well as other nuclear markers, was incompatible with UbiClear^©^. We therefore relied on endogenous AF as an anatomical reference signal.

The arthropod cuticle is coated with a wax layer that protects the insect from environmental exposure. Permeability to water is limited and variable, whereas lipophilic substances penetrate it more easily [27]. We investigated whether dye molecules could pass through the intact arthropod exoskeleton and penetrate deep into the specimen to label internal organs for specific staining with small-molecule dyes and fluorophores. To enhance dye penetration, we used a modified version of the clearing protocol consisting of an initial delipidation step, preceded by a depigmentation and staining step, and finally RI homogenization. This protocol variant was designed to facilitate the diffusion of dyes into whole specimens.

We first tested in woodlice Congo red (CR), a textile dye and classic histological stain that binds β-sheet rich structures like amyloid fibrils and produces red staining. When bound to these ordered structures it shows a red fluorescence [28]. Chitin- and chitosan-rich materials derived from insect and marine shrimp shells exhibit strong Congo red binding capacity [26]. Under basic pH conditions, CR exhibits a crimson coloration, whereas in acidic environments it turns dark blue and functions as a pH indicator [30]. In our experiments, performed at basic pH, CR selectively stained the external cuticle and appendages of the animals in red and it fluoresced in the far red (Fig. 5 A–C).

Similarly, Evans blue (EB) is a protein-binding azo dye commonly used to assess cell viability, membrane integrity, to label neutrophils [31,32], to study vascular permeability and measure blood volume. In mammals, EB strongly binds serum albumin [33] and can act as a vital stain for living cells. When bound to proteins such as serum albumin, its fluorescence increases but it remains weak. Upon 633-nm excitation and with far-red detection, EB fluorescence detection was compatible with lower-wavelength excitation/emission AF imaging, and EB was predominantly detected at the edges of the tergites covering the dorsal plates, as well as small cuticular protrusions (tricorns), thus labeling protein-rich fat body surfaces. In contrast, EB showed little to no staining of the internal organs (Fig. 5D–F).

Finally, Cresyl violet (CV) (commonly used as CV acetate) is a basic aniline dye extensively employed in histology, particularly as a Nissl stain for visualizing neurons [34]. It exhibits strong affinity for acidic cellular components, especially RNA-rich structures. Following depigmentation, specimens were incubated in the post clearing solution. Sequential dual-color light-sheet imaging revealed robust AF under 488-nm excitation with deep red emission from CV when excited at 633 nm [35] (Fig. 5 G–I). In crustaceans, where the ventral nerve cord is located beneath the tergites, the observed staining pattern matched this anatomical arrangement and confirmed the effective penetration of the dye into neural tissue.

### Dual-color autofluorescence and immunofluorescence for multimodal imaging

Given their large size, antibody penetration is expected to occur more slowly and we used the firebug as a case study. An indirect labeling approach with anti-Tuj1 (β-tubulin III) primary antibody and Alexa Fluor 555 secondary antibody was applied (Fig. 6). Although Tuj1 is a common neuronal marker in mammals, its expression in arthropod gonadal tissue makes it suitable for assessing internal labeling. Following clearing, a strong Tuj1 signal was detected, indicating successful antibody penetration into deep tissues. Integration of AF imaging with immunolabeling provided anatomical context and enabled precise localization of the signal. Our results demonstrate that specific immunostaining of cleared arthropods is feasible and compatible with label-free imaging in intact specimens.

## Discussion

### Preserving intact arthropods for volumetric imaging

Optical imaging of intact arthropods remains particularly challenging because their highly pigmented exoskeleton strongly absorbs and scatters light, while the rigid cuticle limits the penetration of clearing agents, dyes, and antibodies. As a result, anatomical investigations often rely on destructive dissection or tissue sectioning, especially for small specimens that are difficult to manipulate. In this study, we show that a rapid tissue-clearing workflow combined with AF imaging, histological staining, and immunolabeling permits volumetric imaging of intact arthropods across diverse taxa and body sizes while preserving their three-dimensional organization.

Formaldehyde fixation proved essential for maintaining tissue integrity throughout clearing and imaging. Although aldehyde fixation has historically been considered problematic in entomology because tissue hardening complicates dissection, this limitation is less relevant in the context of intact volumetric imaging. By stabilizing proteins through methylene bridge formation [36], fixation preserved tissue architecture during depigmentation, delipidation, and refractive-index homogenization while remaining compatible with antibody labeling. Importantly, fixation times were deliberately limited to avoid excessive cross-linking that could impair antibody penetration into the center of the specimen.

### Depigmentation as a major bottleneck in arthropod

Efficient depigmentation remains one of the principal obstacles to arthropod clearing because arthropod pigmentation is chemically diverse and often highly resistant to bleaching. The depigmentation reagent, supplemented with hydrogen peroxide, successfully removed most pigmentation across a broad range of arthropods, including species containing melanin, pteridines and other pigment classes. Depigmentation efficiency increased substantially under basic conditions and elevated temperature, which in turn reduces processing time from weeks to days in some large specimens [37].

Despite these advantages, important limitations remain. Eye pigments, particularly ommochromes [38,39], were frequently more resistant to bleaching than cuticular pigmentation. In addition, crustaceans and other heavily pigmented arthropods occasionally generated gas bubbles during peroxide treatment, likely corresponding to oxygen production [40]. This gas accumulates in the abdomen or appendages of arthropods, which increased the internal pressure, compressing the organs and altering the morphology. These bubbles also produced refractive-index heterogeneities that generated optical artefacts during imaging. Although several approaches have previously been proposed to reduce gas accumulation, including boiling [41] and temperature cycling procedures, bubble formation in appendages remained difficult to eliminate completely, some dissipate over time, yet never fully disappear. The persistence of this phenomenon despite sodium azide treatment, an inhibitor of catalase and peroxidase activity [42], suggests that bubble formation may not result solely from endogenous peroxidase activity, but could instead involve arthropod-specific chemical or enzymatic reactions. Further optimization of depigmentation chemistry will therefore likely be required for large crustaceans and heavily pigmented specimens.

### Rapid clearing and multi-scale imaging across arthropod taxa

Previous approaches for arthropod clearing have relied on potassium hydroxide, sodium hydroxide, lactic acid or pepsin digestion, glycerol, methyl salicylate, or chloral hydrate [43–45], The latter being a sedative, mutagenic and carcinogenic substrate and its use is strictly regulated [46]. While some of these methods improve transparency, they frequently compromise soft-tissue integrity, require prolonged incubation times, or involve hazardous chemicals.

More recently, in 2020, Kuroka applied the CUBIC method to clear large Japanese beetles *Dorcus titanus typhon,* similar in size and coloration to the French *Oryctes nasicornis* shown here [47]. Depigmentation and clearing of these beetles with the CUBIC protocol took 14 and 7 weeks, respectively. By contrast, with the method presented here, the steps require only 17 days and 2 days, respectively, which is a major time advantage for large specimens. Similar to Kuroka, it remains unclear whether fixation fully preserves internal organ integrity, as the size of these insects require a modified light-sheet microscope holder and confocal imaging in not an option here.

The compatibility of the protocol with both confocal and light-sheet microscopy also provides multi-scale imaging flexibility. Light-sheet microscopy allowed rapid volumetric imaging of entire specimens or large body regions with reduced photobleaching and acquisition time, whereas Confocal microscopy offered high-resolution visualization of fine anatomical structures (see e.g., Fig. 5 B–E). However, it should be noted that we used an upright confocal microscope with objectives featuring a long working distance and high numerical aperture.

Because arthropods display strong endogenous AF, imaging can be performed with very low excitation powers in the microwatt range while maintaining high anatomical contrast. In practice, repeated imaging sessions with this reagent did not produce detectable photobleaching, which suggests that cleared specimens can be re-imaged multiple times without significant signal degradation.

### Autofluorescence as an anatomical guide compatible with molecular labeling

AF provided sufficient intrinsic contrast for detailed 3-D visualization of external and internal anatomy without exogenous labeling. (Fig. 3) (Fig. 4). While this fluorescence is clearly multi-component, and different structured are revealed by different excitation wavelength, the relative broad emission spectra make specific AF component detection difficult. Excitation at 405 nm and 488 nm consistently generated the strongest signals, whereas near-infrared excitation produced comparatively weak autofluorescence, thereby opening a spectral window for antibody-based fluorescence detection with limited background interference.

Importantly, AF imaging remained compatible with both histological dyes and immunolabeling (Fig. 5) (Fig. 6). Congo red, previously investigated as a fluorescent stain for crustacean cuticle and Evans blue primarily labelled cuticular structures, while cresyl violet penetrated more deeply and highlighted neural tissues such as the ventral nerve cord. Although these fluorophores are less efficient than modern synthetic dyes such as Alexa Fluor or ATTO derivatives, their lower fluorescence intensity proved advantageous because it remained spectrally compatible with endogenous AF imaging. Furthermore, antibody penetration through intact cleared cuticle was demonstrated using anti-Tuj1 immunolabeling in firebugs, indicating that even relatively large biomolecules can diffuse throughout whole arthropod specimens after clearing.

Together, these observations suggest that AF can serve not only as a label-free imaging modality but also as an anatomical reference framework for targeted molecular labeling approaches. Future developments combining multiplex immunolabeling, spectral imaging, and computational segmentation may therefore facilitate increasingly detailed functional and comparative analyses of arthropod anatomy.

### Toward digital preservation of arthropod biodiversity

Beyond optical transparency, cleared specimens displayed remarkable long-term stability in the post-clearing medium. Specimens stored for several years retained their morphology and remained compatible with repeated imaging. This suggests that the clearing solution may provide an alternative strategy for long-term preservation of delicate arthropod material.

More broadly, volumetric imaging of intact arthropods opens new opportunities for digital preservation of biological collections. Traditional museum preservation methods, like pinning or ethanol storage, remain vulnerable to dehydration, fungal contamination, parasites, and physical degradation over time [48]. In contrast, high-resolution three-dimensional imaging allows the generation of digital replicas that preserve both external morphology and internal anatomy in a non-destructive manner (Supplementary video 1). These datasets can be archived, shared between laboratories, integrated into virtual visualization platforms, and revisited indefinitely without further manipulation of fragile or rare specimens.

Such digital records cannot replace the scientific and historical value of physical collections, but they provide a powerful complementary framework for comparative morphology, morphometric analyses, biodiversity documentation, and long-term accessibility of natural history collections. In this context, tissue clearing combined with volumetric imaging may contribute not only to arthropod anatomy but also to broader efforts aimed at large-scale digitization of biological diversity.

## Conclusion

This work was not intended to describe arthropods, but rather to provide entomologists with a simple, fast, non-toxic and easily accessible pipeline for 3-D imaging of entire specimens. we describe a rapid and broadly applicable workflow for volumetric imaging of intact arthropods that combines tissue clearing, autofluorescence imaging, histological staining, and immunolabeling with standard confocal and light-sheet microscopy. The method preserves three-dimensional anatomy while avoiding destructive dissection and supports long-term preservation and repeated imaging of cleared specimens. Beyond anatomical visualization, this approach establishes a practical foundation for comparative morphology and non-destructive exploration of arthropod biodiversity across research and museum collections.

## Materials and methods

### Sample origin and preparation

A wide variety of arthropods (harlequin bug, grasshopper, mottled shield bug, millipede, wasp, hornet, firebug, fly, spider, cockroach, bee, woodlouse and marine shrimp) were collected in France, euthanized using a commercial pyrethroid-based insecticide and fixed in formalin for 24 h.

### Depigmentation and arthropod clearing

We used an in-house developed, non-toxic, rapid and broadly applicable clearing technique (Delhomme & Oheim, 2023, *Composition and method for clearing biological samples*, patent WO2024089224A1), now commercialized as UbiClear^©^ (Idylle-labs, Paris, France) to clear arthropods in a three-step process. The duration of each step was adjusted according to the size and specific characteristics of the specimen (Table1). Specimens were first bleached using the kit’s depigmentation cocktail, supplemented with hydrogen peroxide (H₂O₂, 3%) at 37–42°C under moderate agitation. Because arthropod heads, legs, and antennae can detach easily, we embedded the depigmented samples in 1% low melt agarose to maintain their integrity during handling and imaging.

Next, arthropods were incubated in the kit’s clearing medium which contains detergents and 2,2-Thiodiethanol (TDE), a chemical reagent that increases the RI, at 37°C under moderate agitation. Finally, cleared specimens were stored in the post-clearing solution (containing 60% TDE and no detergent). Clearing can be reversed when samples are returned to an aqueous buffer with a lower RI (1.33); however, the depigmentation and delipidation are permanent.

### Histologic stains

We performed a series of histological stains on formaldehyde-fixed woodlice using Congo red, Evans blue or Cresyl violet. The protocol was similar for all three dyes, and a variant of the above clearing procedure in which it was split to improve dye penetration. First, delipidation was performed by immersing the fixed woodlice in 1 ml of 0.2 M Tris pH8 solution containing 200 µl of the delipidation solution for 4–5h. This was followed by a 4h depigmentation step. Samples were then incubated overnight in their respective dye solutions:

- Congo red (CAS 573-58-0, *λ* _ex_/ *λ* _em_ = 497 nm/614 nm, RAL Diagnostics) at a concentration of 15 µg/ml.
- Evans Blue (CAS 314-13-6, *λ*_ex_/*λ*_em_ = 620 nm/680 nm, Sigma Aldrich) at a concentration of 200 µg/ml.
- Cresyl violet (CAS 18472-89-4, *λ*_ex_/*λ*_em_= 603 nm/622 nm, Sigma-Aldrich) at 0.001% concentration.

After staining, all specimens went through a series of 30 min rinse cycles, embedded in 1% low-melt agarose, and incubated in post-clearing solution. Samples were kept in this solution until imaging.

### Immunostaining

Similarly, the split version of the clearing protocol was used to improve antibody penetration. We incorporated the delipidation solution into the permeabilization and blocking buffer. Depigmented firebugs were permeabilized and blocked for 5 h in 1 ml of PBS containing 200 µl of the delipidation solution as well as 0.2M Tris buffer pH8, 0.2% Triton X-100, 1% Bovine serum albumin (BSA), 10% goat serum, 10% DMSO, 100 mM glycine and 0.02% sodium azide (NaN₃). The sample was then incubated at 4 °C for 72 hours with the primary antibody anti-Tuj1 (also known as anti-beta tubulin III; GeneTex GTX129913), a pan-neuronal marker in mammals, at a 1:250 dilution in PBS1X supplemented with 5% DMSO, 3% goat serum, 0.001% heparin, and 0.05% NaN₃.

After thorough wash, the sample was incubated overnight with Alexa Fluor 555 Goat anti-Rabbit IgG secondary antibody (Invitrogen) at a 1:500 dilution in the same dilution buffer. Given the long incubation periods, it is important to inhibit the possible microbial or fungi growth with azide. Following additional washes, the firebug was embedded in a low-melting point agarose block and incubated in the usual post-clearing solution. Finally, the sample was stored, still in the post-clearing solution, in the dark until imaging.

### Quantification of tissue clearing

Cleared arthropods were photographed on a macroscope (SMZ800, Nikon Europe BV, Amstelveen, The Netherlands) with a DS-Fi1 camera upon white-light illumination (3000 K). Larger arthropods were photographed with a smartphone (Samsung S24 FE). To evaluate transparency, we placed each specimen on a black-and-white striped target and measured the evolution of Michelson contrast.

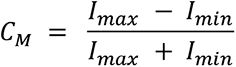

Where C_M_ is the Michelson contrast, I_max_ is the mean intensity of the bright regions, and I_min_ is the mean intensity of the dark regions. Intensities were measured in five regions of interest (ROIs) within the specimen and normalized to the stripe intensities outside the tissue using Fiji/ImageJ software [49]. Contrast was also measured in a sample-free background region and the final transparency was expressed as a percentage of the background contrast, with 0% representing a completely opaque specimen and 100% a fully transparent one.

### Multimodal fluorescence imaging

#### Macro-imager

For multi-wavelength overview imaging, we used a macro imager (Sapphire FL, Azure Biosystems, Sierra Court, Dublin, CA) that offers up to 5-μm lateral resolution. Images with different combinations of excitation and emission wavelengths were taken to generate an AF excitation-emission matrix.

#### Confocal microscopy

Small specimens were mounted in 3-D printed sample holders, made from black polylactic acid (PLA), with cavity depths ranging from 1–10 mm. The cavities were filled with the post-clearing reagent and covered with a standard coverslip (BK-7, Marienfeld Superior, Braunschweig, Germany) and sealed with fast-setting dental silicone (Twinsil speed 22 silicon, Rotec, France). Confocal micrographs were acquired on an upright confocal laser-scanning microscope (CLSM) (LSM710 META, ZEISS, Oberkochen, Germany) fitted with a 10x/NA0.45 air objective (working distance, *wd.* = 2 mm) using Zen Black 2.1 software (Zeiss). AF images were captured upon 405-nm excitation with 47µW (2%) laser power and detection in the 410- to 740-nm band for broad AF collection. The pinhole diameter was systematically set to 1 Airy unit (AU).

#### Light-sheet microscopy

Larger samples were imaged using a custom-built light-sheet microscope [50]; (H. Touja, B. Chauvin *et al., in preparation*) featuring a compact dual-arm illuminator with cylindrical beam-shaping optics and a central sample holder. Light-sheet positioning was controlled manually in *x*, *y* and via a motorized precision stage (Physik Instrumente) in *z*, while the entire module moved in x,y relative to the detection path on a precision motorized stage (Märzhäuser, Wetzlar, Germany). Illumination was provided by a fiber-coupled four-line laser combiner (445, 488, 561, and 638 nm; Oxxius), generating ∼3 µm-thick light sheets (at waist) that were co-aligned and intensity-matched, and used either simultaneously or sequentially with subsequent image fusion (custom blending or alpha-blending in ImageJ). Samples were mounted in optical glass cuvettes or custom holders and imaged in the post clearing solution. Fluorescence was collected on an upright macroscope (MVX10, EVIDENT Olympus Europe, Hamburg, Germany) equipped with long-working-distance objectives (1×/NA0.25 air, *wd.* = 65 mm, or 2×/NA0.5 air, *wd.* = 20 mm) and a motorized focus drive (Märzhäuser). Imaging parameters included *z*-steps of 2–6 µm, laser powers of 1–50 µW, and exposure times of ∼ 400 ms - 1.2 s per plane for label-free imaging, depending on the specimen. For most experiments, we used a multi-band Quad405/488/561/640 filter set (AHF Analysentechnick, Tübingen, Germany), permitting simultaneous multi-line excitation for AF imaging or sequential acquisition for labelled samples; in some cases, single-band filter sets (Olympus U-MGFP/XL, U-MRFPHQ/XL). Cresyl violet fluorescence was excited at 561 nm and detected through a 595/50H band-pass filter (AHF).

#### Stitching

Due to illumination geometry, the effective homogeneous imaging region of the light sheet was limited to an approximately cuboidal volume of ∼2 cm along the propagation axis, 0.8–1 cm orthogonal to the propagation axis, and ∼20 µm in thickness. Larger specimens were therefore imaged as overlapping stacks that were subsequently stitched together computationally.

### Image processing

Images were processed using IMAGEJ/FIJI software. 3-D rendering and animations were performed with IMARIS (Bitplane, Oxford Instruments, Zürich, Switzerland).

## Acknowledgements

We acknowledge the core facility of BioMedTech Facilities Université Paris Cité INSERM US36 | CNRS UAR2009 | Université Paris Cité for assistance with custom mechanical pieces and microscopy and Brieuc Chauvin (SPPIN) for help with light-sheet microscopy. Hervé Suaudeau and Luc Tamisier (SPPIN) provided the hardware and software architecture for large-scale data acquisitions, storage and backup. La Pitancerie, the shared garden of Cachan (Val de Marne, France) deserves a special mention for its insect biodiversity and was a valuable site for specimen collection. The bees were obtained from Luc Tamisier’s hive in Charenton, (Valde-Marne, France).

## Author contributions

MM: Methodology, Data curation, Investigation, Sample preparation, Confocal acquisitions, Data analysis, Writing – original draft. HT: Light-sheet acquisitions. BD: Conceptualization, Methodology, Investigation, Writing – original draft. MO: Funding acquisition, Project administration, Supervision, final manuscript writing.

## Funding Declaration

This work was supported by the French Centre National de la Recherche Scientifique (CNRS) and by grants from the French Agence Nationale de la Recherche (ANR-23-ce19-0006-01 KIARA), from the Université Paris Cité (IDEX Emergence KLEYA), FranceBioImaging (a large-scale national infrastructure initiative, FBI, ANR-10-INSB-04, Investments for the future) and by the Région Île-de-France via a DIM C brains equipment grant (M-Cube). None of these organizations had any influence on the design or outcome of this st udy.

## Data availability statement

The data that support the findings of this study are available from the corresponding author upon reasonable request.

## Abbreviations

AF: Autofluorescence
AU: Airy unit
Conf /: CLSM Confocal laser-scanning microscopy
LS /: LSFM Light-sheet fluorescence microscopy
NIR: Near-infrared
NA: Numerical aperture
3-D: Three-dimensional
2-D: Two-dimensional
TDE: 2,2′-Thiodiethanol
H₂O₂: Hydrogen peroxide
RI: Refractive index
BSA: Bovine serum albumin
NaN₃: Sodium azide
DMSO: Dimethyl sulfoxide
Tris: Tris(hydroxymethyl)aminomethane buffer
PBS: Phosphate-buffered saline
CR: Congo red
EB: Evans blue
CV: Cresyl violet
VG: Venom gland
ST: Stinger
MA: Mandible
MXP: Maxillary palpus
GL: Glossa
LP: Labial palpus
EY: Eye
AN: Antenna
NADA: N-acetyldopamine
NBAD: N-β-alanyldopamine
ROI: Region of interest
CM: Michelson contrast
I_max_ / I_min_: Maximum / minimum intensity
λ_ex_ / λ_em_: Excitation / emission wavelength

## Additional Information

### Competing interest statement

the authors declare no competing interests. BD and MO are co-inventors of patent WO2024089224A1, entitled ‘Composition and method for clearing biological samples’ owned by the Centre National de la Recherche Scientifique (CNRS) and licensed for commercialization by SATT ErgaNeo. The clearing pipeline, developed in our laboratory, is now marketed under the brand name UbiClear© by Idylle-Labs (Paris, France) www.idylle-labs.com. This work was entirely performed at the SPPIN (CNRS/University Paris-Cité), and no company or funder had any role in the study design, experiments or outcome. The authors have no financial interest in publishing this method.

---

**Supplementary figure 1:**
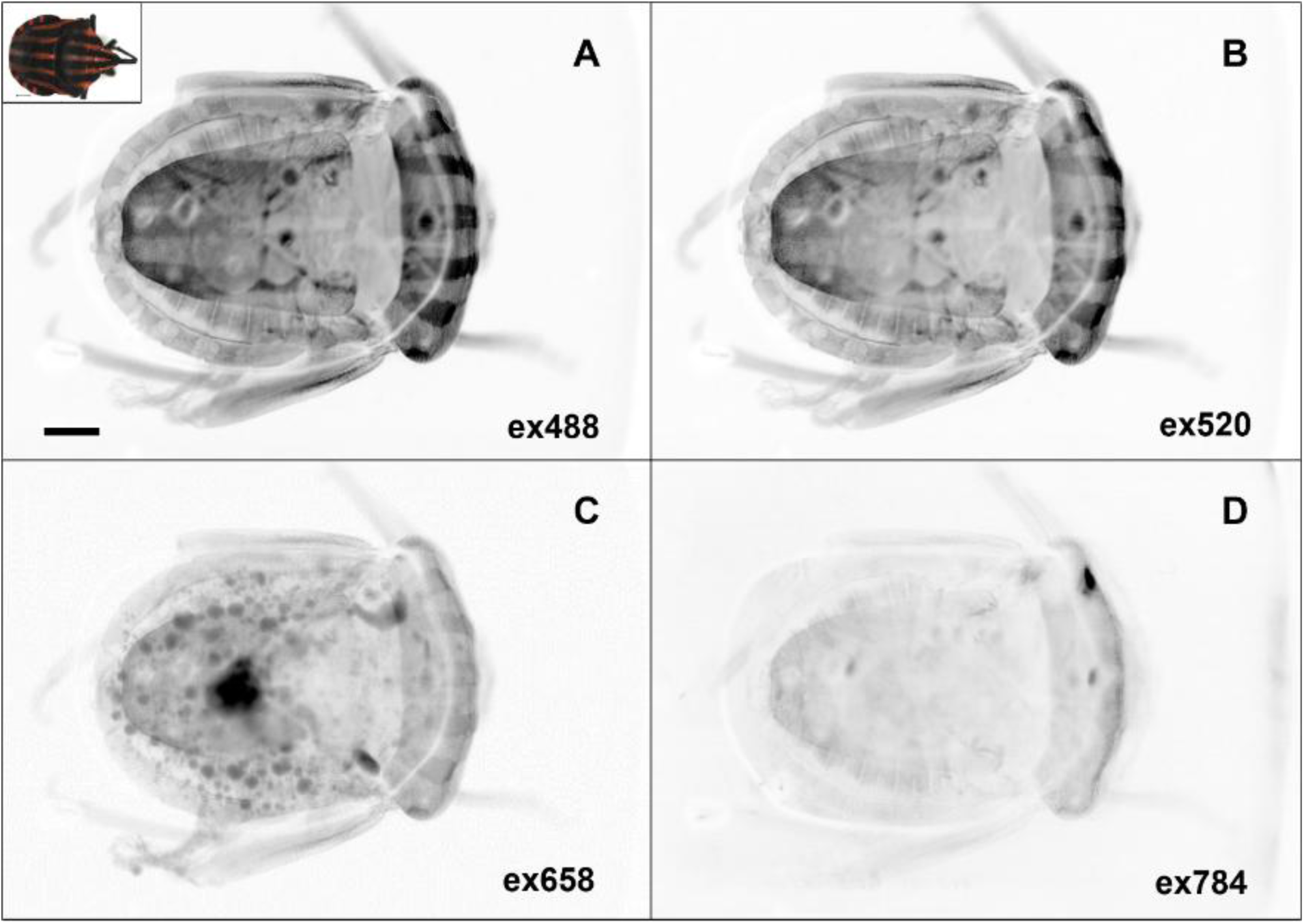
Comparison of autofluorescence under different excitation wavelengths. Two-dimensional images of a harlequin bug *(Graphosoma lineatum)* acquired with a Sapphire Imager (Azure Biosystems) under white-light illumination before and after UbiClear^©^ processing. AF was recorded at multiple excitation wavelengths: *(**A**)* 488 nm, *(**B**)* 520 nm, *(**C**)* 658 nm and *(**D**)* 784 nm. Scale bar: 1 mm.

*Supplementary Video 1:* Confocal AF imaging of a cleared wasp at 10x magnification acquired using 405 nm excitation. The z-stack’s total depth is 1300μm with a z-step of 100μm. Scale bar: 1mm.

## Notes

### Competing Interest Statement

The authors have declared no competing interest.

